# DARPP-32 in motor cortex regulates structural and synaptic plasticity in corticothalamic neurons and enables motor learning

**DOI:** 10.64898/2026.05.01.722248

**Authors:** Pisanò Clarissa Anna, Aaltonen Alina, Spanu Valeria, Tamaki Ayu, Fisone Gilberto, Santini Emanuela, Borgkvist Anders

**Affiliations:** Department of Neuroscience, Karolinska Institutet, 17177 Stockholm Sweden

## Abstract

The dopamine and cAMP-regulated phosphoprotein of 32 kDa (DARPP-32), a key mediator of monoaminergic signaling, is expressed in the cortex; however, its cellular distribution and role in cortically dependent behaviors remain elusive. Here, we determined the functional integration of DARPP-32 in motor cortex circuitry using molecular profiling, circuit tracing, patch-clamp electrophysiology, virus-assisted gene targeting and motor behavior analyses. Unlike the significant overlap between DARPP-32 and dopamine receptors in striatal GABAergic medium spiny projection neurons, we found that the majority of DARPP-32-positive cortical neurons express the corticothalamic marker FoxP2 but not dopamine D1 or D2 receptors. Notably, in cortical slices, adenylyl cyclase activation induced a more robust increase in DARPP-32 phosphorylation at threonine 34, a protein kinase A target site, compared to dopamine D1 receptor stimulation. Conditional ablation of DARPP-32 in the motor cortex did not affect basal or psychostimulant-induced motor activity but reduced motor aptitude and compromised overnight retention of motor skill. Concomitantly, the absence of DARPP-32 reduced dendritic spines density and prevented the induction of glutamatergic long-term potentiation in layer 6 motor cortical neurons. Altogether, our study demonstrates a critical role for DARPP-32 in cortical synaptic plasticity, emphasizing its importance in corticothalamic regulation of motor skill learning.

## Introduction

Motor learning enables organisms to flexibly adapt their behavior to changing environmental demands. In mammals, the motor cortex serves as a critical substrate for the acquisition, storage, and retrieval of motor skills (Peters, Liu, & Komiyama, 2017). During learning, cortical circuits engaged in the behavior undergo synaptic remodeling, resulting in the formation of new synapses and the elimination of unused connections (Economo, Komiyama, Kubota, & Schiller, 2024). This structural reorganization enhances neuronal communication and progressively expands the cortical representation of the learned behavior, culminating in the refinement of a motor map (Kida & Mitsushima, 2018; Monfils, Plautz, & Kleim, 2005).

Although the motor cortex is essential for motor skill acquisition, the specific cell types and molecular mechanisms that mediate this process remain incompletely understood. Accumulating evidence identifies dopamine as a key modulator of motor learning (Cousineau, Plateau, Baufreton, & Le Bon-Jégo, 2022; C. Vitrac & Benoit-Marand, 2017). In rodents, disruption of dopaminergic input to the motor cortex impairs the acquisition of new motor skills (Hosp, Pekanovic, Rioult-Pedotti, & Luft, 2011). These deficits are accompanied by reduced long-term synaptic plasticity (Molina-Luna et al., 2009; Rioult-Pedotti, Pekanovic, Atiemo, Marshall, & Luft, 2015), impaired dendritic spine remodeling (Guo et al., 2015), and abnormal motor map formation (Hosp, Molina-Luna, Hertler, Atiemo, & Luft, 2009), indicating that dopamine supports cortical restructuring during learning. These findings are clinically relevant, as impaired motor learning is a hallmark of neurological conditions such as stroke and Parkinson’s disease (Clément Vitrac, Nallet-Khosrofian, Iijima, Rioult-Pedotti, & Luft, 2022) (Chu et al., 2024), both of which involve compromised dopaminergic signaling. Understanding the molecular mechanisms linking dopamine to cortical plasticity may therefore provide new strategies to enhance rehabilitation and promote functional recovery.

The dopamine- and cAMP-regulated phosphoprotein of 32 kDa (DARPP-32) is a central mediator of dopamine receptor signaling (Greengard, 2001). When phosphorylated by second-messenger pathways downstream of dopamine receptors, DARPP-32 potently inhibits protein phosphatase 1 (PP1). Because PP1 regulates the phosphorylation state of ion channels, kinases, and transcription factors, DARPP-32 functions as a key integrator of dopaminergic and glutamatergic inputs, thereby influencing neuronal excitability and synaptic plasticity (Onn, Fienberg, & Grace, 2003) (Bateup et al., 2010; Calabresi et al., 2000; Fienberg et al., 1998). Genetic inactivation of DARPP-32 in mice impairs dopamine-related behaviors (Bateup et al., 2010; Fienberg et al., 1998; Nairn et al., 2004) and disrupts motor learning (Qian, Forssberg, & Diaz Heijtz, 2015), consistent with a critical role in determining dopamine-dependent adaptations.

However, the role of DARPP-32 in the motor cortex remains unresolved. Most studies focused on the striatum, where DARPP-32 is highly expressed and plays a pivotal role in motor control. Notably, DARPP-32 is also expressed in cortical neurons (Ouimet, Miller, Hemmings, Walaas, & Greengard, 1984) and it has been implicated in the dopaminergic modulation of cognitive and executive functions (Berger, Febvret, Greengard, & Goldman-Rakic, 1990). In rodents, DARPP-32 signaling mediates dopaminergic control of long-term synaptic plasticity in the frontal cortex and object recognition memory (Hotte et al., 2007; Hotte et al., 2006). Reduced cortical DARPP-32 expression has been reported in patients with schizophrenia (Devor et al., 2017), a disorder characterized by dopaminergic dysfunction and cognitive impairment. Despite these observations, dopaminergic signaling in the cortex differs substantially from that in the striatum. Cortical dopamine innervation is comparatively sparse, and DARPP-32 may engage distinct cellular and molecular mechanisms. For instance, dopamine regulation of cortical interneurons can occur independently of DARPP-32 (Trantham-Davidson, Kröner, & Seamans, 2008), and receptor expression analyses indicate that dopamine receptors are enriched in non-pyramidal intracortical neurons (Anastasiades, Boada, & Carter, 2019), whereas DARPP-32 is preferentially expressed in pyramidal neurons projecting to the thalamus (Ouimet, 1991). Thus, it remains unclear whether DARPP-32 regulates synaptic plasticity and motor learning in the motor cortex, and in which neuronal population it exerts its effects.

To address these questions, we combined retrograde tracing and immunostaining with transgenic reporter mice to map DARPP-32 expression in relation to D1- and D2-expressing neurons. To determine whether cortical DARPP-32 is required for behavior, we performed virus-assisted genetic inactivation and assessed motor activity induced by novelty and amphetamine, two paradigms sensitive to DARPP-32 function. Finally, we tested whether structural and synaptic plasticity underlying motor skill learning depend on cortical DARPP-32 by quantifying dendritic spine density, long-term synaptic plasticity, and motor performance in the rotarod task. Our results identify a critical role of cortically expressed DARPP-32 in motor learning, indicating that its function in the cortex, as opposed to the striatum, supports circuit-specific regulation of motor behavior.

## Results

### Distribution and circuit organization of DARPP-32-expressing neurons in the motor cortex

Previous anatomical studies of the prefrontal and cingulate cortex have reported DARPP-32 expression in neurons located in layers 5 and 6 (L5, L6), which comprise major output layers projecting to the contralateral cortex, striatum, thalamus and spinal cord (Kuroiwa et al., 2012; Ouimet, LaMantia, Goldman-Rakic, Rakic, & Greengard, 1992; Ouimet et al., 1984; Walaas & Greengard, 1984). To determine whether DARPP-32 exhibits a similar distribution in the motor cortex, we co-stained brain sections with antibodies against DARPP-32 and the vesicular glutamate transporter 2 (vGlut2), a marker of thalamocortical input layers (Graziano, Liu, Murray, & Jones, 2008; Herzog et al., 2001; Hisano et al., 2000) (Fig 1A, B). Fluorescent cells were quantified in confocal images from the primary (M1, MOp) and secondary motor cortex (M2, MO), spanning the region from the appearance of the forceps minor rostrally to the caudal midline fusion of the corpus callosum, thereby encompassing the rostral motor cortex as defined by recent anatomical classification (Chon, Vanselow, Cheng, & Kim, 2019). Consistent with previous findings in frontal cortex, DARPP-32 positive cells were enriched in L6, defined here as the region between the corpus callosum and the vGluT2-stained L5b (Fig 1B).

**Figure 1.**
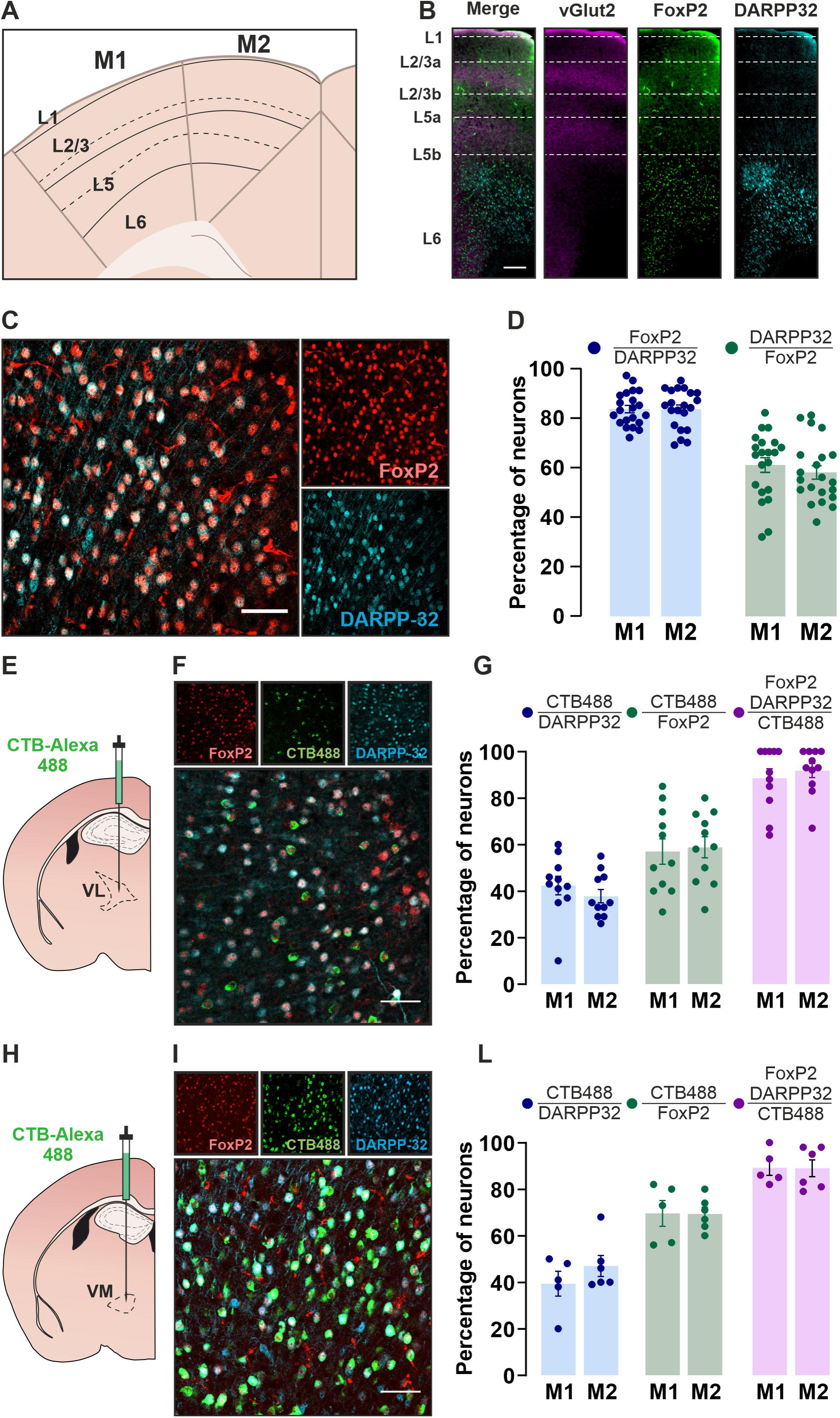
DARPP-32 is expressed in corticothalamic neurons innervating ventrolateral and ventromedial thalamus. **(A)** Schematic of primary (M1) and secondary (M2) motor cortex with cortical layers delineated. **(B)** Representative confocal images showing immunostaining for VGluT2 (magenta), FoxP2 (green) and DARPP-32 (light blue) across the vertical cortical column from pia (top) to white matter (bottom). Scale bar = 25 mm **(C)** Representative confocal images showing immunostaining for DARPP-32 (blue) and FoxP2 (red) in M2. Scale bar = 50 µm **(D)** Bar graphs showing the fraction of FoxP2+ DARPP-32-expressing neurons (left) and DARPP-32+ FoxP2-expressing neurons (right) in M1 and M2. (n=21 slices, 6 mice). **(E)** Schematic of fluorescent retrotracer (CTB488) injection into the ventrolateral (VL) thalamus. **(F)** Representative confocal images showing immunostaining for FoxP2 and DARPP-32 along with CTB488 in the MCtx of mice injected with CTB488 in VL thalamus. **(G)** Bar graphs showing the fraction of DARPP-32+ (D32, blue) and FoxP2+ (green) neurons in the M1 and M2 that are retrogradely labelled from VL with CTB488, and the fraction of CTB488-labelled neurons expressing DARPP-32, FoxP2, or both markers. Mean±SEM, n=11 slices, 4 mice. Scale bar = 50 µm. **(H)** Schematic of CTB488 injection into the ventrolateral (VM) thalamus. **(I)** Representative confocal images showing immunostaining for FoxP2 and DARPP-32 along with CTB488 in the MCtx of mice injected with CTB488 in VM thalamus. Scale bar = 50 µm. **(J)** Bar graphs showing the fraction of DARPP-32+ (D32, blue) and FoxP2+ (green) neurons in the M1 and M2 that are retrogradely labelled from VM with CTB488, and the fraction of CTB488-labelled neuron expressing DARPP-32, FoxP2, or both markers. Mean±SEM, n=5-6 slices, 2 mice. Scale bar = 50 µm.

**Figure 1.**
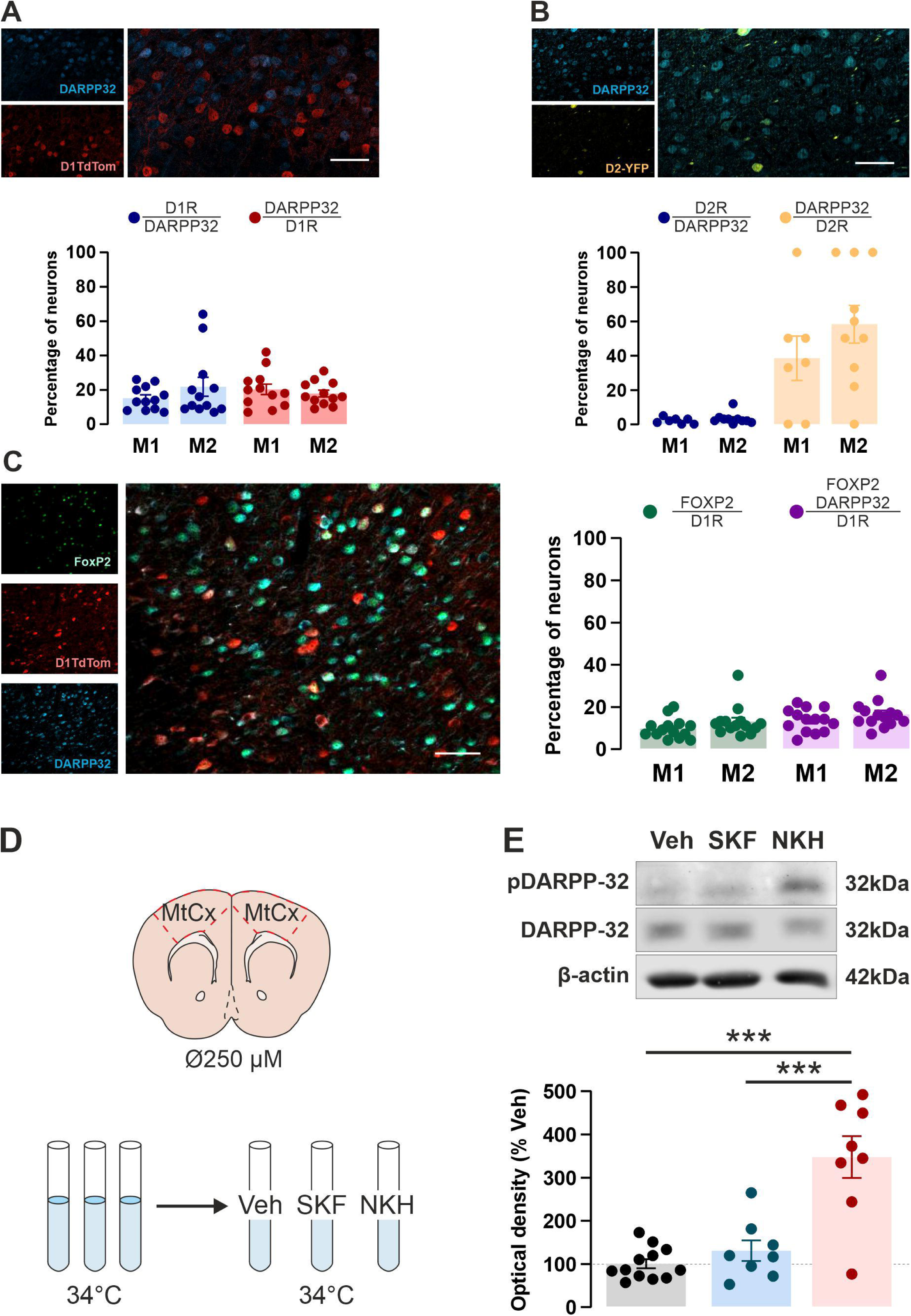
Anatomical and functional segregation of DARPP-32-expressing neurons from putative D1 and D2 receptor-expressing cells in the MCtx. **(A)** Representative confocal images (upper panel) showing immunostaining for DARPP-32 (blue) and tdTomato (red, tdTom) in Drd1a-tdTomato mice. Scale bar = 50 µm. Bar graph (bottom panel) showing the fraction of DARPP-32+ neurons expressing tdTom (blue bars) and tdTom+ neurons expressing DARPP-32 (red bars) in M1 and M2. Mean±SEM, n=14 slices, 4 mice. **(B)** Representative confocal images (upper panel) showing immunostaining for DARPP-32 (blue) and expression of eYFP driven by D2-Cre (Drd2-Cre mice, AAV5-DIO-YFP, yellow, eYFP+). Scale bar = 50 µm. Bar graphs (bottom panel) showing the fraction of DARPP-32+ neurons expressing eYFP (blue bars) and eYFP+ neurons expressing DARPP-32 (yellow bars) in L6 M1 and M2. Mean±SEM, n=7-10 slices, 4 mice. **(C)** Representative confocal images (left panel) showing immunostaining for DARPP-32 (blue) and tdTom (red) and FoxP2 (green) in Drd1a-tdTomato mice. Scale bar = 50 µm. Bar graphs (right panel) showing the fraction of tdTom+ neurons expressing FoxP2 (green bars), or FoxP2 and DARPP-32+ (magenta bars) in M1 and M2. (n=14 slices/group, 4 mice). **(D)** Schematic of the brain slice pharmacology experiment. **(E)** Representative immunoblots (upper panel) of pThr34-Darpp-32, total DARPP-32, used as dissection control, and actin employed as loading control. Lower panel shows the quantification of pThr34-Darpp-32. 1-way ANOVA, F (2, 24) = 20.98, ***P<0.001, followed Tukey post-hoc. veh vs skf ns; veh vs nkh ***P>0.001; skf vs nkh ***p<0.001. Mean±SEM, n=8-12 slices/group, 3 mice.

This layer-restricted expression pattern suggests that DARPP-32-positive cells are glutamatergic pyramidal neurons, characterized by dendrites extending toward superficial layers and axons projecting to either subcortical or intracortical targets (Briggs, 2010). To determine which of these neuronal subtypes express DARPP-32, we performed co-immunostaining using antibodies against DARPP-32 and Forkhead Box P2 (FoxP2), a transcription factor selectively expressed in glutamatergic corticothalamic neurons (Qi et al., 2025) (Fig. 1B). Quantification of single- and double-labeled neurons revealed a strong overlap between DARPP-32 and FoxP2 expression, indicating that DARPP-32 is enriched in corticothalamic neurons (Fig 1C, D). However, not all FoxP2 neurons expressed DARPP-32 (Fig 1D) suggesting that DARPP-32 is restricted to a subpopulation of glutamatergic corticothalamic neurons.

To identify thalamic targets of DARPP-32-positive L6 corticothalamic neurons, we injected fluorescently labeled cholera toxin subunit B (CTB) into the ventrolateral (VL) and ventromedial (VM) thalamic nuclei (Fig 1E, H), two recipients of motor cortical projections (Muñoz-Castañeda et al., 2021). We then imaged CTB labeled cells in the motor cortex of slices stained for DARPP-32 and FoxP2 (Fig 1F, I). Because this approach labels only a fraction of target-specific neurons, the number of CTB-labeled FoxP2 neurons was taken as an estimate of the maximal retrogradely labeled population at each thalamic injection site. CTB injection into the VM labeled 38 ± 6% (mean ± SEM, n = 5-6 slices from 2 mice) of FoxP2 neurons in both M1 and M2, whereas VL injection labeled 23 ± 3% (mean ± SEM, n=11 slices from 4 mice), consistent with denser motor cortical innervation of the VM compared to the VL (Muñoz-Castañeda et al., 2021). In both VM and VL injected mice, fewer CTB-labeled neurons expressed DARPP-32 compared to FoxP2 (Fig 1G, L). Importantly, the majority of CTB-labeled DARPP-32-positive neurons co-expressed FoxP2 (Fig 1G, L). These results indicate that DARPP-32 is expressed in a subpopulation of corticothalamic projection neurons across different thalamic targets.

Taken together, these anatomical analyses demonstrate that DARPP-32 is expressed in a subset of corticothalamic projection neurons in the motor cortex, mirroring expression patterns previously described in the prefrontal and cingulate cortex.

### DARPP-32 expression in relation to dopamine D1 and D2 receptors

DARPP-32 is highly expressed in striatal medium spiny neurons (MSNs), where it co-localizes with dopamine receptors and regulates their downstream signaling. In the rodent cortex, L6 receives the densest meso-cortical dopaminergic input and contains neurons expressing dopamine D1 and D2 receptors (Yger & Girault, 2011), raising the possibility that DARPP-32 may similarly regulate dopaminergic signaling in this region. To assess whether DARPP-32 is co-expressed with D1 and D2 receptors, we stained brain slices from transgenic mice carrying fluorescent proteins expressed under transcriptional control of the promoter of D1 or D2 receptor, using DARPP-32 antibodies (Fig 2A-D).

**Figure 2.**
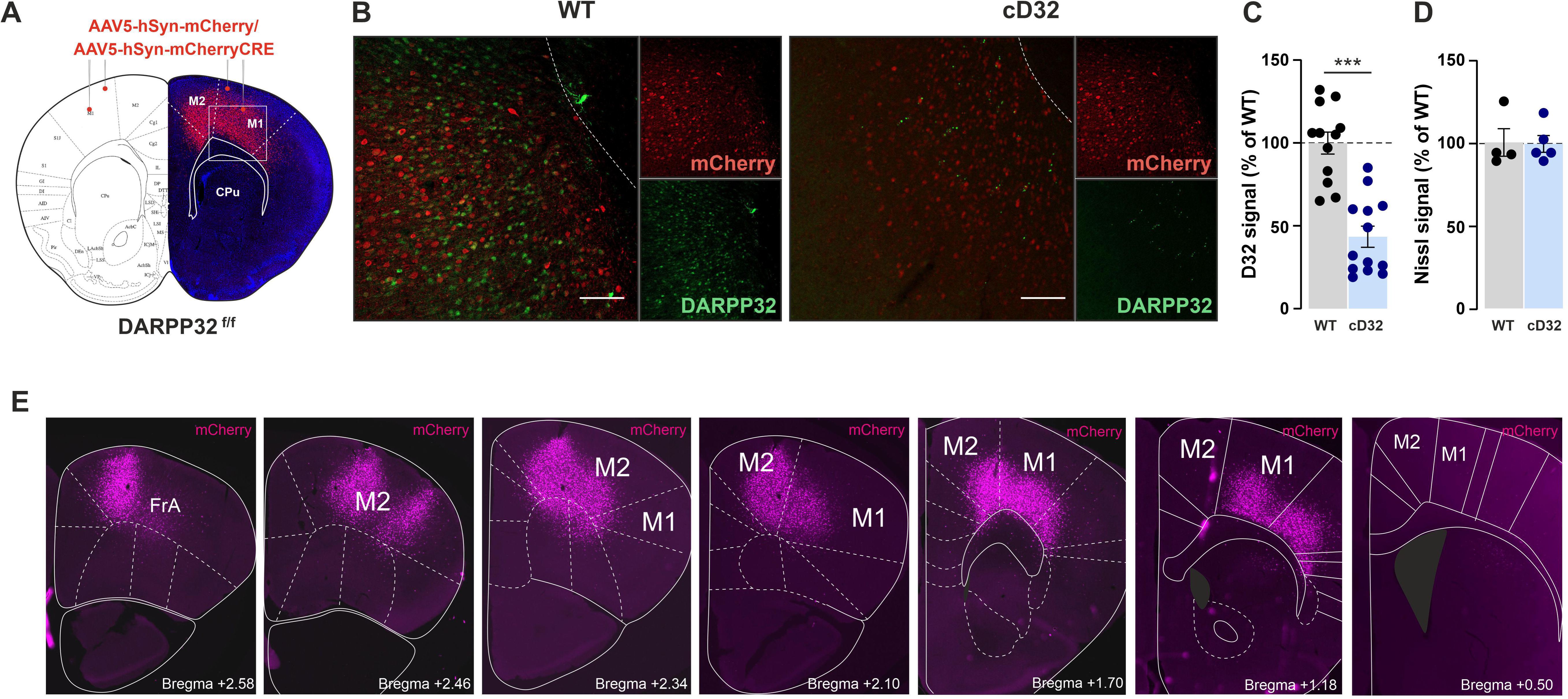
AAV-assisted ablation of DARPP-32 from neurons in the MCtx. (A) Schematic of the targeted region (adapted from Paxinos brain atlas) and a representative confocal image from a DARPP-32^f/f^ mouse bilaterally infused with AAV-hSyn-Cre-mCherry (cD32) in MCtx, illustrating mCherry expression. Control mice consisted of DARPP-32^f/f^ littermates injected in the same region with AAV-hSyn-mCherry (WT). Nuclei are counterstained with DAPI (blue). (B) Representative confocal images showing DARPP-32 immunofluorescence (green) and mCherry (red) in M1 of WT and cD32 mice. Dashed line marks the dorsal border of the white matter. Scale = 100 µm (C) DARPP-32 immunofluorescence in mCherry-expressing regions of M1/M2 in WT and cD32 mice. Data represent mean±SEM normalized to WT signal intensity. Unpaired t-test. t(23)=6,145, ***p<0.001 (n=12-13 mice per group). (D) Nissl signal in mCherry-expressing regions of M1/M2 in WT and cD32 mice. (E) Representative mCherry fluorescence in M1/M2 of a cD32 mouse. Outlined subregions and bregma coordinates according to Paxinos (Paxinos, 2001).

In Drd1a-tdTomato mice (Shuen, Chen, Gloss, & Calakos, 2008), 15±2% and 22±6% of DARPP-32-positive neurons in primary (M1) and secondary (M2) motor cortex, respectively, co-expressed tdTomato. In addition, only 20±3% and 18±2% of the tdTomato-positive neurons in the M1 and M2 expressed DARPP-32 (Fig 2A, B), a markedly lower fraction compared to the striatum where virtually all D1 receptor-expressing MSNs co-express DARPP-32 (Valjent, Bertran-Gonzalez, Hervé, Fisone, & Girault, 2009).

In Drd2-Cre transgenic mice injected with an AAV transducing Cre-dependent YFP, 2±7% (M1) and 3±1% (M2) of DARPP-32-positive neurons were YFP-positive (Fig 2C, D). However, among YFP-expressing neurons, 39±13% (M1) and 58±11% (M2) expressed DARPP-32, suggesting a greater overlap of DARPP-32 with D2 than D1 receptors.

Previous studies have shown that D1 receptors are present in intratelencephalic neurons of the prefrontal cortex, but not in subcortically projecting neurons (Anastasiades et al., 2019). To determine whether D1 receptor-positive neurons expressing DARPP-32 in the motor cortex belong to corticothalamic populations, we performed co-immunostaining on slices from Drd1-tdTomato mice using antibodies against DARPP-32 and the corticothalamic marker FoxP2. This analysis revealed a small percentage of D1 receptor-positive neurons co-expressing DARPP-32 and FoxP2 (Fig 2E, F).

To test whether DARPP-32 in the motor cortex mediates dopamine receptor signaling, we treated cortical slices with either the D1 receptor agonist SKF81297 or adenylyl cyclase activator NKH-477. Both compounds increase the canonical protein kinase A-mediated phosphorylation of DARPP-32 at Thr34 in the striatum (Svenningsson et al., 2004). Using western blot analysis with phosphorylation-state-specific DARPP-32 antibodies, we found that SKF81297 (5 min, 1 µM) did not significantly increase phospho-Thr34-DARPP-32 (Fig 2H). In contrast, NKH-477 (5 min, 10 µM) significantly increased phospho-Thr34-DARPP-32 (Fig 2I). These results indicate that in motor cortex, D1 receptors are functionally uncoupled from cAMP/PKA-mediated DARPP-32 phosphorylation, likely because DARPP-32 is predominantly expressed in corticothalamic neurons that lack D1 receptors.

Altogether, our findings suggest that most DARPP-32-expressing neurons in the motor cortex do not carry D1 or D2 receptors, revealing a notable difference from the expression pattern in dopamine receptor-expressing MSNs.

### DARPP-32 ablation in motor cortex does not affect novelty-induced exploration or amphetamine-induced hyperactivity

Genetic inactivation of DARPP-32 reduces novelty-induced exploration and psychostimulant-induced hyperactivity (Bateup et al., 2010; Bonito-Oliva, DuPont, Madjid, Ögren, & Fisone, 2016; Fienberg & Greengard, 2000). To assess whether DARPP-32 in the motor cortex contributes to these behaviors, we conditionally inactivated DARPP-32 by injecting an AAV-Cre-mCherry into the motor cortex of DARPP-32^fl/fl^ mice (cD32; Fig 3A). Littermates receiving AAV-mCherry injections served as controls (WT). Five weeks post-surgery, DARPP-32 immunostaining was significantly reduced in regions expressing Cre-dependent mCherry fluorescence (Fig 3B-C), while Nissl staining intensity, reflecting neuronal density, was indistinguishable between cD32 and WT mice (Fig 3D).

**Figure 3.**
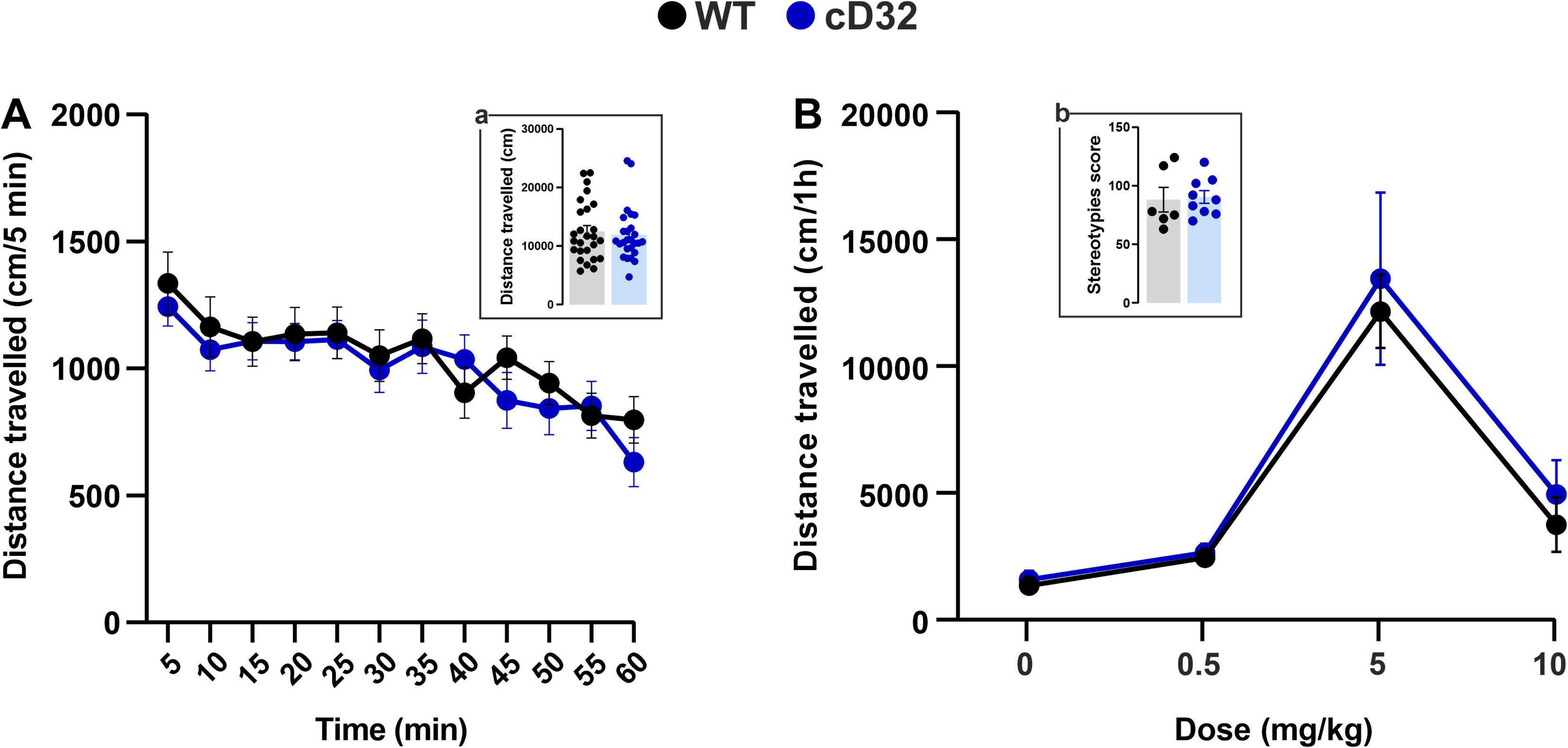
DARPP-32 ablation in motor cortex does not alter exploratory behavior or acute response to the psychostimulat amphetamine. **(A)** Distance travelled across time in WT (black) and cD32 (blue) mice exposed to a novel cage enivornment. n=26 mice per group. Two-way RM ANOVA showed a significant effect of Time F (7,433, 371,7) = 14,27, ***p<0.001, no signficant effect of genotype F (1, 50) = 0,1976, p=0.6586, and no significant time x genotype interaction F (11, 550) = 1.040, P=0.4089. Bonferroni post-hoc. (**a**) Total distance travelled. (n=26 mice per group). **(B)** Distance travelled following amphetamine injections. Two-way ANOVA showed a signficant effect of dose F (3, 46) = 36.67, ***p<0.001, no signficant effect of genotype F (1, 46) = 0.7712, p=0.3844, and no significant time x genotype interaction F (3, 46) = 0.1389, p=0.9362. **(b)** Combined stereotypies score following injection of high dose amphetamine (10 mg/kg, i.p.). n=5-12 mice per group. All data in figure represent mean ±SEM.

When introduced to a novel cage environment, both WT and cD32 mice displayed a gradual decrease in locomotor activity over time (Fig 4A). Habituation to novelty, reflected by this progressive decline, as well as total distance traveled over the 1-hour session, did not differ between genotypes (Fig 4A a). Systemic amphetamine administration induced comparable dose-dependent locomotor activation in WT and cD32 mice (Fig 4B). In both groups, locomotion peaked at 5 mg/kg, while 10 mg/kg produced stereotypies that reduced the locomotion relative to 5 mg/kg. The severity of stereotypies was also similar between WT and cD32 mice (Fig 4B).

**Figure 4.**
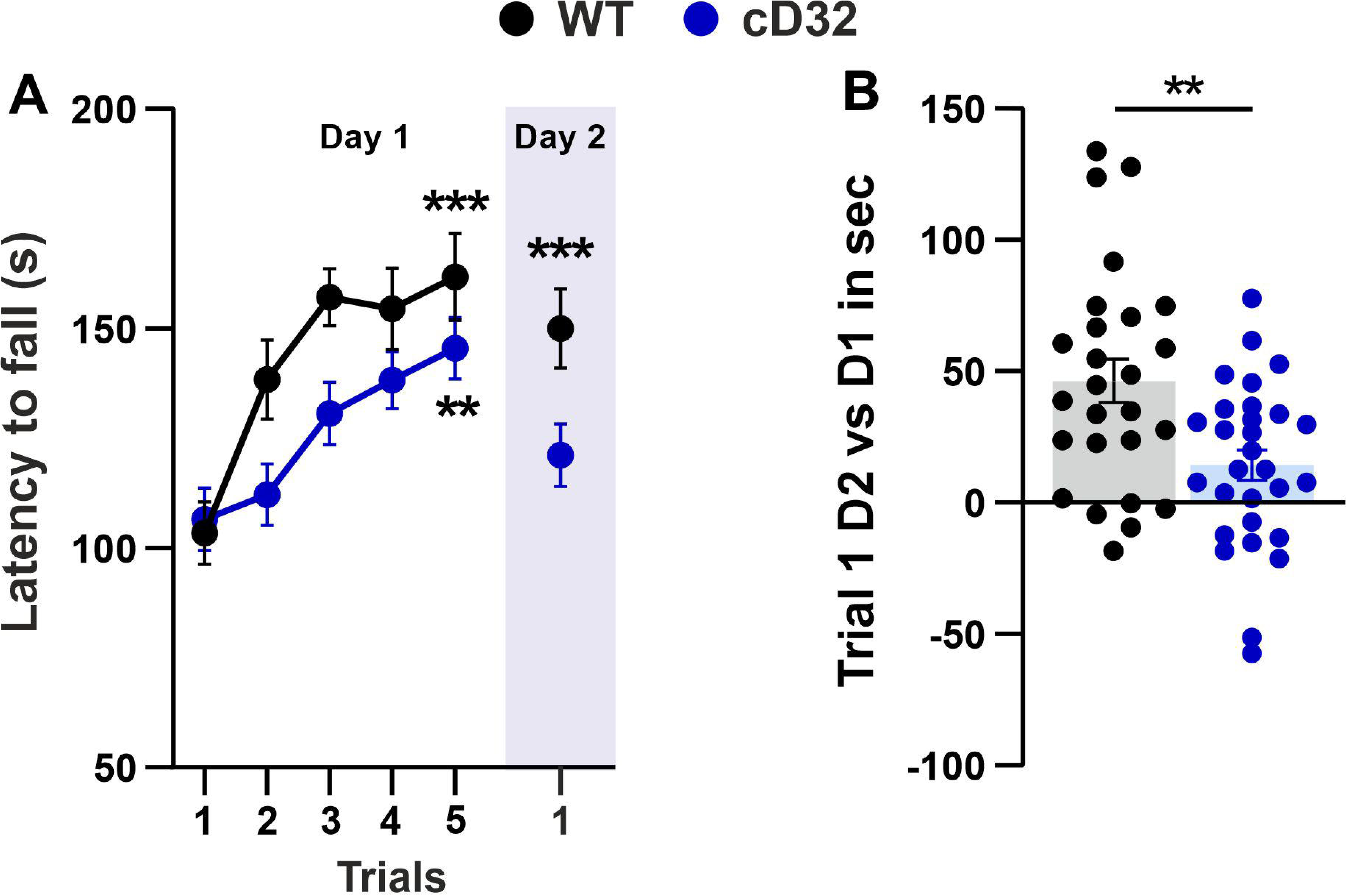
Cortical DARPP-32 ablation impairs motor performance and memory consolidation in the accelerated rotarod test. **(A)** Latency to fall across five trials on day one and during one re-exposure trial on day two. Two-way RM ANOVA showed a significant effect of time F (4.120, 218.4) = 17.78, ***p<0.001, genotype F (1, 53) = 5.467, *p<0.05 and no time x genotype interaction F (5, 265) = 2.021, p=0.0760. n = 26-29 mice. ***p<0.001, **p<0.01, *p<0.05 different from Trial 1, Bonferroni post hoc comparison. **(B)** Difference in latency to fall between the re-exposure trial on day two and the first trial on day one. t-te*st* t(53)=3.23. **p<0.01 WT vs. cD32. All data in figure represent mean ±SEM.

These results indicate that DARPP-32 in motor cortex does not contribute to novelty-induced exploration or amphetamine-induced hyperactivity, in contrast to the effects observed following global DARPP-32 inactivation.

### DARPP-32 ablation in the motor cortex impairs motor learning

Loss-of-function mutation in DARPP-32 leads to motor skill learning deficits (Qian et al., 2015). To investigate whether these deficits reflect impaired DARPP-32 signaling in the motor cortex, we subjected cD32 and WT mice to the accelerated rotarod task, which assesses motor performance, including balance, coordination, and motor skill acquisition (Costa, Cohen, & Nicolelis, 2004). Task performance was quantified as the latency to fall across five consecutive training trials on day one. To assess consolidation of skill memory, the mice received a single re-exposure trial on day two.

WT and cD32 mice showed similar latency to fall on the first training trial, indicating that DARPP-32 inactivation in motor cortex does not impair baseline motor function. However, although performance improved in both groups with training, cD32 mice consistently exhibited shorter fall latencies across trials, indicating a critical role for DARPP-32 expression in the motor cortex for motor skill acquisition (Fig 5A).

**Figure 5.**
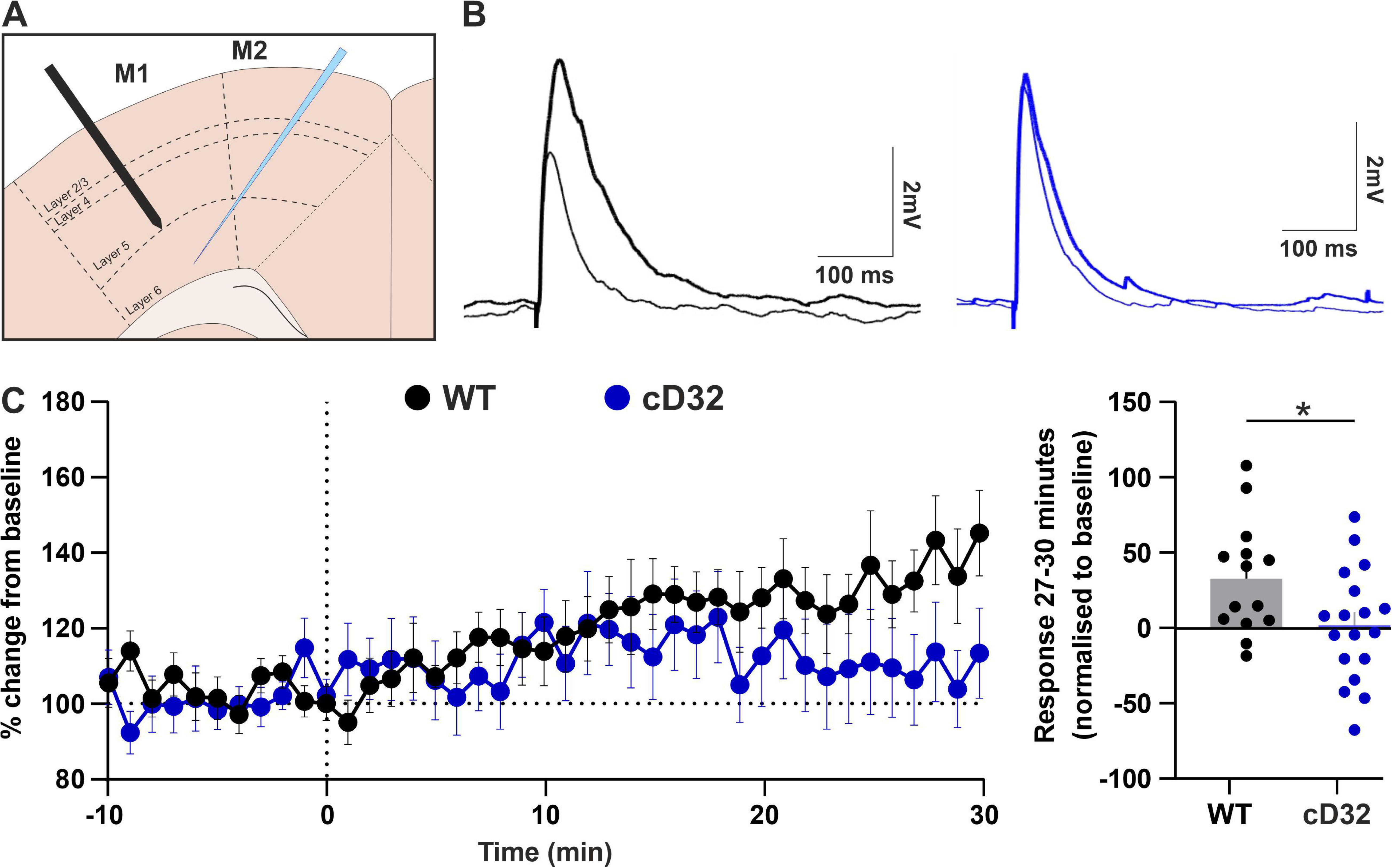
Cortical ablation of DARPP-32 prevents induction of long-term potentiation in L6 M1 neurons. **(A)** Schematic representation of the regions of electrical field stimulation (black electrode) and whole-cell patch clamp recording (blue). **(B)** Representative evoked excitatory synaptic potentials (EPSPs) recorded in mCherry fluorescent cells of L6 in the MCtx of slices from WT (black traces) and cD32 (blue traces) mice. Thin traces and thick traces represent before and after LTP induction by theta burst stimulation, respectively. **(C)** Time course of EPSP amplitude expressed as a percentage of baseline. Dashed line marks the time of theta burst stimulation. **(D)** Percentage increase in EPSP amplitude measured 27-30 minutes after stimulation. t-test. *t*(30)=2.33, *p<0.05. All data in figure represent mean ±SEM.

Upon re-exposure on day two, WT mice maintained performance levels comparable to their final training trial, whereas cD32 mice showed a significant decline in fall latency (Fig 5A). To directly compare performance between WT and cD32 mice, we calculated the change in fall latency between the first training trial and the re-exposure trial. WT mice exhibited significantly greater performance gains than cD32 mice (Fig 5B).

Together, these findings demonstrate that expression of DARPP-32 in the motor cortex is essential for optimal motor skill acquisition and next-day retention, consistent with a role in motor skill consolidation.

### DARPP-32 ablation in motor cortex impairs induction of long-term potentiation

Motor skill learning depends on the strengthening of glutamatergic synapses in motor cortex (Monfils & Teskey, 2004; Yang et al., 2009). To test whether the learning deficits observed in cD32 mice reflect impaired long-term synaptic plasticity, we recorded picrotoxin-resistant excitatory postsynaptic potentials (EPSPs) from mCherry-positive L6 neurons in acute motor cortex slices. EPSPs were elicited by electrical stimulation of L5b to target superficial thalamocortical synapses (Fig 6A). Baseline EPSP amplitudes were similar in WT and cD32 mice (WT = 4.1 ± 0.3 mV vs. cD32 = 4.0 ± 0.3 mV, t(30) = 0.288, p>0.05, t-test, n = 14-18 cells, 8 mice per group), indicating intact basal glutamatergic synaptic transmission.

**Figure 6.**
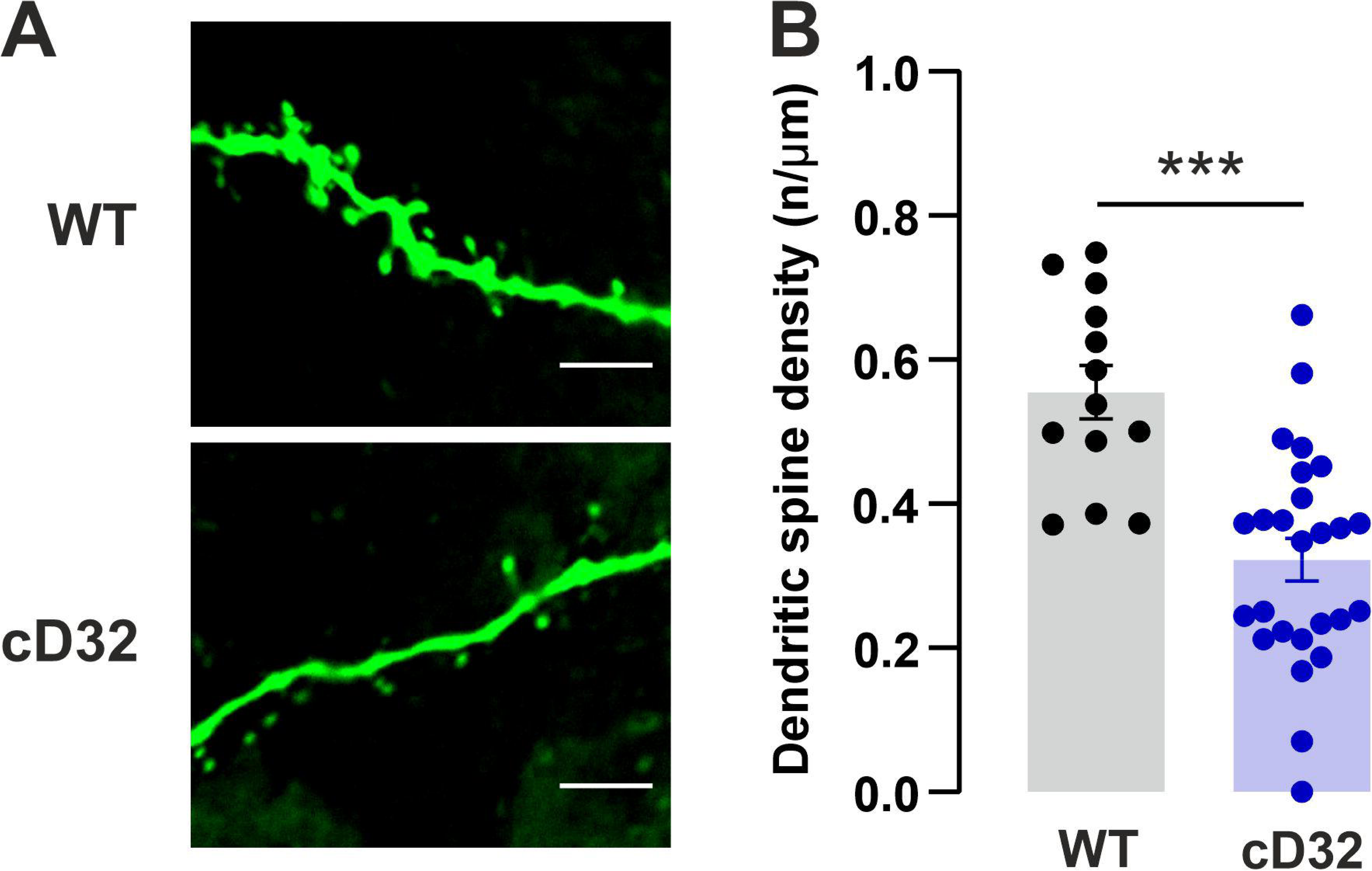
Cortical DARPP-32 ablation results in reduced spine density in L6 M1 neurons. **(A)** Representative confocal images of Lucifer Yellow-filled dendritic segments of L6 neurons in the MCtx of slices from WT (top) and cD32 (bottom) mice. Scale bar = 5 µm. **(B)** Bar graph showing dendritic spine density in WT and cD32 L6 neurons. Data are presented as mean ±SEM. t-test, t(37) = 4.714, ***p<0.001, n =13-26.

Theta burst stimulation, which induces long-term potentiation (LTP) in L2/3 (Hess, Aizenman, & Donoghue, 1996), produced a robust increase in EPSP amplitude in WT slices (Fig 6B-C). In contrast, no potentiation was observed in cD32 slices (Fig 6C). Thirty minutes after theta stimuli, EPSP amplitudes were significantly smaller in neurons from cD32 mice compared to WT (Fig 6D). These results suggest that motor learning impairments in cD32 mice reflect reduced capability for synaptic plasticity at glutamatergic synapses of L6 neurons.

### DARPP-32 ablation in the motor cortex reduces spine density in L6 neurons

Both motor skill learning and LTP are associated with increased dendritic spine density in motor cortex (Engert & Bonhoeffer, 1999). To determine whether the behavioral and synaptic deficits observed in cD32 mice are accompanied by alteration in dendritic spine density, we filled L6 cortical neurons in motor cortex slices from WT and cD32 mice with Lucifer Yellow (Dumitriu, Rodriguez, & Morrison, 2011) and imaged dendritic segments using confocal microscopy (Fig 7A).

Quantification of dendritic spines on distal apical and basal dendrites (∼100 µm from the soma) revealed a significant reduction in spine density in cD32 compared to WT neurons (Fig 7B), consistent with impaired dendritic spine maintenance or turnover in the absence of DARPP-32.

## Discussion

In this study, we identified DARPP-32 as a critical regulator of activity-dependent plasticity in a sub-population of corticothalamic neurons in the motor cortex and demonstrate that its expression is required for the consolidation of motor skill learning.

DARPP-32 has been extensively characterized in the striatum, where it is selectively expressed in GABAergic MSNs and plays a central role in mediating dopaminergic signaling underlying motor behaviors. However, in cortical areas, DARPP-32 has been reported in glutamatergic projection neurons rather than inhibitory populations (Kuroiwa et al., 2012; Ouimet et al., 1984; Perez & Lewis, 1992). This circuit-dependent cell-type specificity suggests a distinct role for DARPP-32 in modulating excitatory transmission and cortical information processing.

To define the cellular substrates of DARPP-32 signaling in the motor cortex, we examined its expression across genetically defined neuronal populations. DARPP-32 was enriched in deep-layer neurons and predominantly expressed in corticothalamic projection neurons, as indicated by co-expression with the corticothalamic transcription factor FoxP2 (Ferland, Cherry, Preware, Morrisey, & Walsh, 2003; Fong, Kuo, Wu, Chen, & Liu, 2018; Hisaoka, Nakamura, Senba, & Morikawa, 2010; Sorensen et al., 2015). This overlap is consistent with a role for FoxP2 in regulating DARPP-32 expression, potentially via direct binding to the promoter region of the DARPP-32 gene (*Ppp1r1b*) (Hisaoka et al., 2010; Vernes et al., 2011). Accordingly, previous studies show that FoxP2 perturbation reduces DARPP-32 expression in both cortex and striatum (Co, Hickey, Kulkarni, Harper, & Konopka, 2020).

DARPP-32 expression was largely restricted to a subset of FoxP2-positive neurons, consistent with enrichment in corticothalamic populations. Retrograde tracing studies revealed these neurons project to ventrolateral and ventromedial thalamus. However, it is likely that additional DARPP-32-expressing neurons also target subregions of the motor thalamus not examined in this study. This possibility is supported by anatomical studies showing that FoxP2-positive corticothalamic neurons in other cortical regions, including mouse somatosensory cortex (PMID: 3986047), innervate multiple thalamic nuclei spanning both first- and higher-order regions. In the motor thalamus, the ventrolateral and ventromedial nuclei are considered first-order and higher-order, respectively (Bosch-Bouju, Hyland, & Parr-Brownlie, 2013), indicating a similar organizational logic.

Overall, our findings demonstrate projection-specific enrichment of cortical DARPP-32, distinguishing it from the homogeneous expression in striatal MSNs, and indicate that DARPP-32 signaling in the cortex is organized along defined output pathways.

### DARPP-32 and dopamine receptors are segregated in the motor cortex

Our study reveals a marked segregation between DARPP-32 and dopamine receptor-expressing neurons in the motor cortex. In line with previous work in prefrontal cortex (Cieslak et al., 2024), D1 receptor-expressing neurons were present in deep motor cortical layers. However, only a small subset co-expressed DARPP-32 and the corticothalamic marker FoxP2. Instead, the majority of DARPP-32-positive neurons were associated with corticothalamic populations, whereas D1 receptor-expressing neurons more likely correspond to intratelencephalic projection neurons or interneurons, as reported in other cortical areas (Anastasiades et al., 2019; Co et al., 2020; Gaspar, Bloch, & Le Moine, 1995; Hisaoka et al., 2010; Le Moine & Gaspar, 1998; Trantham-Davidson et al., 2008). This interpretation is further supported by evidence that DARPP-32 does not mediate D1-dependent modulation of excitability in the prefrontal cortex (Trantham-Davidson et al., 2008), suggesting that DARPP-32 operates through distinct signaling pathways in cortical circuits.

The segregation of DARPP-32 and D1 receptor-expressing neurons in the motor cortex is also supported by a paralleled functional dissociation. Pharmacological stimulation of D1 receptors in acute brain slices produced little increase in DARPP-32 phosphorylation at Thr34, whereas direct activation of adenylyl cyclase robustly increased phospho-Thr34-DARPP-32, indicating that DARPP-32 in the motor cortex remains responsive to cAMP/PKA signaling but is not strongly coupled to D1 receptor activation. Although a small subset of neurons may exhibit D1-dependent DARPP-32 phosphorylation below the detection threshold, the overall lack of responsiveness supports a limited functional coupling between D1 receptors and DARPP-32 in the motor cortex.

Similar to striatal MSNs, D1 and D2 receptors are largely segregated across neuronal populations in the cerebral cortex (Cieslak et al., 2024; Wei et al., 2018). This organization prompted us to examine whether DARPP-32 is preferentially co-expressed with D2 receptors. We observed a comparable segregation between DARPP-32 and D2 receptors, although a greater fraction of D2 receptor positive neurons co-expressed DARPP-32 than D1 receptors.

This distribution aligns with the known neurochemical profile of D2 receptor-expressing neurons in the rodent cerebral cortex, which is distinct from that of DARPP-32. D2 receptors are found in GABAergic interneurons as well as in corticostriatal and corticocortical projection neurons, but are principally absent from corticothalamic neurons (Ferrari, 1979; Yu et al., 2019; Zhang et al., 2021). In contrast, DARPP-32 does not appear to be expressed in cortical GABAergic interneurons (Kuroiwa et al., 2012), and is instead enriched in corticothalamic neurons, as indicated by our findings. While DARPP-32 expression was more prevalent among D2-expressing cortical neurons than D1-expressing neurons, the absolute number of D2 positive neurons was exceedingly low. Thus, despite this enrichment, the small population size suggests that DARPP-32 signaling in D2 neurons is unlikely to exert a substantial influence on global dopaminergic modulation of motor cortical circuits.

Together, these results reinforce the conclusion that, in contrast to its canonical role in the striatum, DARPP-32 in the motor cortex operates largely independently of classical dopaminergic signaling pathways. Rather, DARPP-32 is preferentially expressed in corticothalamic neurons, where it remains responsive to cAMP/PKA signaling and can couple intracellular signaling to activity-dependent plasticity in cortical circuits underlying motor learning.

### Cortical DARPP-32 is required for motor learning

The present study established cortical DARPP-32 as a critical regulator of motor learning. Selective reduction of DARPP-32 expression in the motor cortex of adult mice impaired performance on the accelerating rotarod, particularly during re-exposure, indicating a deficit in motor skill consolidation rather than initial acquisition.

These findings position DARPP-32 as a key molecular component supporting the persistence of motor memories and highlight a phase-specific cAMP/PKA/DARPP-32 signaling across brain regions. While previous work has implicated striatal DARPP-32 in early acquisition phases of motor skill (Qian et al., 2015), our results demonstrate that cortical DARPP-32 is required for consolidation. Together, these observations support a model in which DARPP-32 signaling is distributed across circuits in a temporally structured manner, with engagement shifting from striatal circuits during acquisition to cortical circuits during consolidation.

Additional evidence supports a role for DARPP-32 in corticothalamic-dependent motor learning. FoxP2 mutant mice, which exhibit deficits in corticothalamic neuron function, display impaired performance in motor learning tasks such as the accelerating rotarod (Groszer et al., 2008). Given the enrichment of DARPP-32 in FoxP2-positive corticothalamic neurons, these observations are consistent with a contribution of this neuronal population to motor skill learning, although a direct mechanistic link remains to be established.

Similarly, global DARPP-32 knockout mice exhibit deficits in motor coordination and performance in rotarod tasks, consistent with disrupted motor circuit function (Jeljeli, Strazielle, Caston, & Lalonde, 2003; Sakayori et al., 2019). These impairments likely reflect altered activity within corticothalamic loops, which are essential for the precise timing and coordination of motor output. Importantly, in our study, selective reduction of DARPP-32 in the motor cortex did not affect novelty-induced exploration or amphetamine-induced locomotion, indicating that these behaviors are primarily mediated by striatal circuits (Bateup et al., 2010; Bonito-Oliva et al., 2016).

Together, these findings indicate that DARPP-32 in the motor cortex specifically contributes to motor skill consolidation via corticothalamic circuits, rather than to general locomotor activity or dopamine-dependent behavioral responses.

### DARPP-32 links synaptic and structural plasticity to motor learning in corticothalamic circuits

The behavioral deficits observed following cortical DARPP-32 ablation are paralleled by impairments in both synaptic and structural plasticity. Electrophysiological recordings revealed a marked reduction in LTP in the motor cortex, indicating that DARPP-32 is required for activity-dependent strengthening of glutamatergic synapses. This aligns with studies in the striatum, where genetic deletion of DARPP-32 abolishes both LTP and long-term depression (LTD) (Bateup et al., 2010; Calabresi et al., 2000), underscoring its conserved role as a key regulator of synaptic plasticity across brain regions.

Notably, the mechanisms by which DARPP-32 supports plasticity in the cortex appear to differ from those in the striatum. Dopamine-dependent plasticity in the motor cortex has been shown to rely on phospholipase C-dependent signaling pathways that bypass PKA (Rioult-Pedotti et al., 2015), suggesting that DARPP-32 does not mediate dopamine-driven synaptic modulation in this region. Instead, our findings suggest that DARPP-32 contributes to cortical plasticity independently of canonical dopaminergic signaling.

In addition to functional plasticity, DARPP-32 depletion caused a significant reduction in dendritic spine density, indicating disruption of structural synaptic substrates. This effect is consistent with the involvement of DARPP-32 in cytoskeletal regulation through its interaction with adducin, a key modulator of actin dynamics in dendritic spines (Engmann et al., 2015). Together, these findings demonstrate that DARPP-32 coordinates both functional and structural plasticity necessary for stabilizing motor memories.

At the circuit level, our findings suggest that DARPP-32-dependent plasticity operates within parallel pathways that converge on the thalamus. In the basal ganglia, DARPP-32 in striatal MSNs regulates motor output indirectly via thalamic modulation, whereas in the cortex, DARPP-32 in corticothalamic neurons provides a direct route for shaping thalamic activity. The convergence of these pathways onto shared thalamic targets, such as the ventrolateral and ventromedial nuclei, points to a coordinated mechanism through which DARPP-32-dependent plasticity across circuits contributes to motor learning (Ramot et al., 2025).

Alterations in cortical DARPP-32 expression have been implicated in neuropsychiatric disorders, including schizophrenia and psychosis (Kunii et al., 2011; Nishiura et al., 2011), and have been linked to behavioral responses to antipsychotic treatment (Chana et al., 2013; Kuroiwa et al., 2012). Within this framework, our findings highlight DARPP-32 as a critical molecular node connecting synaptic plasticity, circuit function, and behavior across brain regions.

## Concluding model and implications

While it has long been known that the motor cortex plays a critical role in motor learning, the precise cellular circuit components involved have only recently begun to be explored in detail using molecular mapping and circuit manipulation approaches. Recent studies have demonstrated that corticothalamic projections regulate motor skill learning (Li et al., 2026) and may do so by inhibiting corticospinal neurons (Carmona et al., 2025).

Our findings support a model in which DARPP-32 operates as a circuit-specific regulator of plasticity across parallel motor pathways. While in the striatum, DARPP-32 integrates dopaminergic signals to shape action selection and motor output, we show that in the motor cortex it is preferentially expressed in corticothalamic neurons, where it regulates activity-dependent synaptic and structural plasticity required for motor learning. These parallel DARPP-32-dependent mechanisms converge on shared thalamic targets, providing a potential substrate for coordinated plasticity across cortico-basal ganglia-thalamic loops.

More broadly, our results indicate that DARPP-32 in the motor cortex functions beyond its canonical role as a mediator of dopaminergic signaling, linking cAMP signaling to activity-dependent synaptic plasticity and circuit adaptation. By dissociating dopamine-dependent and dopamine-independent functions of DARPP-32 across brain regions, this work provides a framework for understanding how common molecular signaling pathways are repurposed in distinct neuronal populations to support different phases of learning.

## Limitations and future directions

Although our anatomical and functional data indicate a segregation between DARPP-32 and dopamine receptor signaling in the motor cortex, we cannot fully exclude that a small subset of neurons exhibits D1-dependent DARPP-32 activation below the detection threshold of our biochemical assays. Second, while our findings demonstrate that DARPP-32 remains responsive to cAMP/PKA signaling, the upstream sources of cAMP in corticothalamic neurons remain unidentified. These may include neuromodulatory inputs or activity-dependent intracellular pathways, which were not directly addressed in the present study. Third, our viral approach targeted DARPP-32 in deep cortical layers but did not resolve potential cell-type-specific contributions within corticothalamic subpopulations. Future studies combining cell-type-specific manipulations with in vivo circuit recordings will be necessary to determine how DARPP-32-dependent plasticity shapes corticothalamic output during motor learning.

## Acknowledgements

We thank members of Borgkvist, Santini and Fisone laboratories for technical support and fruitful scientific discussions. The behavior analyses were conducted at the Animal Behavioral Core Facility (ABCF) at Karolinska Institutet. We are grateful for veterinary guidance and excellent animal service provided by the staff in the Department of Comparative Medicine. This work was supported by Swedish Research Council grants 2016-03129_VR (AB), 2024-03684_VR (A.B), 2016-02758_VR (E.S), 2023-02943_VR (E.S) and 2024-02966_VR (G.F), R-2024, Strategiska forskningsområdet neurovetenskap (StratNeuro) at Karolinska Institutet (A.B and E.S), Parkinsonfonden 1543/24 (A.B), Åhlén-stiftelsen mA4h18 (A.B), Magnus Bergvalls Stiftelse 2018-02764 (A.B), Olle Engkvist Stiftelse 207-0575 (A.B), Knut och Alice Wallenbergs Stiftelse grants 2017-0169 (E.S) and 2020-0054 (E.S).

## Materials and Methods

### Animals

C57BL/6NRJ mice (20-30g, 2.5-3.5 months old) used for tracing experiments were obtained from Jackson Laboratory. Drd1-tdTomato (Shuen et al., 2008), DARPP-32 f/f (Bateup et al., 2010), and D2Cre mice were taken from our inbred colonies.

The animals were housed in groups, up to six mice, in individually ventilated cages, under standardized conditions (12 h light/dark cycle, stable temperature 22°C, and humidity 40–50%) with food and water ad libitum. All experiments were carried out in accordance with the guidelines of Research Ethics Committee of Karolinska Institutet and Swedish Animal Welfare Agency. All efforts were made to minimize animal suffering and to reduce the number of animals used.

### Stereotactic Surgery

Mice were anesthetized with isoflurane (4%, maintained at 1.5-2% during surgery) and placed in a stereotaxic frame with a heating pad to maintain normothermia. Each received 0.1 mg/kg Temgesic (buprenorphine) subcutaneously at the start of surgery. Anesthesia depth was monitored by checking pedal and palpebral reflexes, muscle tone, and respiration. Ophthalmic ointment was applied to prevent corneal drying. The surgical site was shaved, disinfected, and treated with local anesthetic (Xylocaine), followed by a midline incision (∼1 cm). The skin was retracted, the skull cleaned, and stereotactic coordinates were zeroed on bregma. Head position was verified by moving the injection needle along the sagittal suture, and adjustments were made if needed. A small hole was drilled at the injection site coordinates (Paxinos, 2001).

#### For retrograde tracing

100 nL of CTB AlexaFluor 488 was injected into the VM/VL (AP: −1.46, ML: 0.8, DV: −4.25; AP: −1.10, ML: 1.10, DV: −3.28) at 0.2 µL/min.

#### For D2 mapping

a 1:1 mixture of AAV5-EF1a-DIO-eYFP and AAV5-hSyn-mCherry (100 nL) was injected into three sites of the rostral motor cortex (AP: 1.34, ML: −1.25/-1.5/-1.9, DV: −0.8/-1/-1.2) at 50 nL/min.

#### For DARPP-32 conditional KO

mice received two injections per hemisphere of 100 nL AAV5-hSyn-mCherry-cre (or AAV5-hSyn-mCherry for controls) at (AP: +1.34, ML: ±1.2, DV: −0.8) and (AP: +1.34, ML: ±1.90, DV: −1.20).

After injections, the needle remained for 5 minutes to ensure proper delivery before being withdrawn. The site was disinfected, the incision closed with stitches, and the mice were returned to their cages once fully recovered.

### Tissue Fixation and Histology

Mice were anesthetized with pentobarbital (60 mg/kg, i.p) and perfused transcardially with 4% paraformaldehyde (PFA) in 0.1 M sodium phosphate buffer, pH 7.4. Brains were post-fixed overnight (for immunofluorescence) or 1 hour (for Lucifer Yellow filling) in the same solution at 4°C, then stored in Phosphate-buffered saline (PBS) until they were processed (Lieberman et al., 2020). Brains from CTB-injected mice were sectioned at 30-40 μm using a VT1000s vibratome (Leica Biosystems) and stored at −20°C in 0.1 M sodium phosphate buffer with 30% ethyleneglycol and 30% glycerol. Brain regions were identified using a mouse brain atlas (Paxinos, 2001). Free-floating sections from the CTB or LY experiment were rinsed three or six times for 10 min in TBS, incubated for 1 hour in 10% Normal Goat Serum (NGS), 0.6% Triton X-100 in TBS, and rinsed three times. They were then incubated overnight at 4°C with primary antibodies (Table 1). Sections were rinsed three times for 10 min in TBS and incubated for 30 min with Alexa Fluor secondary antibodies (1:500), washed in PBS, and mounted with Fluoromount-G™ with or without DAPI (depending on the channel used). LY-filled sections were incubated overnight with Streptavidin Alexa 488 (1:500, 2% NGS, 1% Triton X-100 in TBS). Images were obtained using a Zeiss LSM 800 Airyscan confocal microscope (20X-40X objective, 2.5x digital zoom) with consistent laser and detector settings. Image analysis was performed in ImageJ.

### Dendritic spine reconstruction

Corticostriatal coronal slices (150 µm) were obtained using a VT1000s vibratome (Leica Biosystems). Lucifer Yellow CH (Invitrogen) was dissolved at 8% in 50 mM Tris HCl. The filling protocol was adapted from Dumitrius et al. (Dumitriu et al., 2011). Micropipettes, pulled from borosilicate glass capillaries (OD 1.5 mm, ID 0.86 mm, 10 cm length), had a resistance of 40-50 MΩ when filled with Lucifer Yellow. Neuronal fillings were performed using an electrophysiology rig with an Axopatch 200B amplifier (Molecular Devices), 10x dry and 40x water-immersion lenses (Olympus), VX55 camera (TILL Photonics), and Solis LED fluorescence system (Thorlabs). RFP-positive neurons in layer 6 (M1 and M2) were filled. LY was injected incrementally with −2 nA for 5 minutes, −5 nA for 5 minutes, and −10 nA for 30 seconds, totaling 10.5 minutes. Leaking neurons were discarded.

### Analysis of dendritic spines density

ImageJ was used for dendritic spine image analysis. Image stacks of dendritic sections were analyzed frame by frame. The counter tool was used to mark spines and the segmented line function was used to measure the length of the dendritic section. The spine density was calculated by dividing the number of spines by the length of the dendritic section.

### Acute Brain Slice

Acute brain slices were prepared as described by Lieberman et al. (2018)(Lieberman et al., 2018). Briefly, mice underwent cervical dislocation, and the brain was quickly placed in ice-cold high sucrose cutting solution (in mM): 10 NaCl, 2.5 KCl, 25 NaHCO3, 0.5 CaCl2, 7 MgCl2, 1.25 NaH2PO4, 180 sucrose, 10 glucose, bubbled with 95% O2/5% CO2 (pH 7.4). Brains were sectioned into 250 μm coronal slices (including the striatum) using a VT1200 vibratome (Leica Biosystems). Slices were transferred to ice-cold cutting solution, and the motor cortex was dissected. Slices were then placed in ACSF (in mM): 125 NaCl, 2.5 KCl, 25 NaHCO3, 2 CaCl2, 1 MgCl2, 1.25 NaH2PO4, 10 glucose, bubbled with 95% O2/5% CO2 (pH 7.4) at 34°C. Slices from one mouse were split into three conditions (Vehicle, SKF-38393 1μM, NKH-477 Forskolin 10μM) and rested for 1 hour. After 5 minutes of drug incubation at 34°C, slices were homogenized by sonication in 1% SDS.

### Western Blot

Equivalent protein amounts (25-40 μg per sample) were loaded into 10% gradient polyacrylamide gels, following Santini et al. 2007 (Santini et al., 2007). Proteins were transferred to Immobilon FL PVDF membranes (0.2 μm pore size). Blots were blocked in PBST with LiCor medium for 1 hour at room temperature, then incubated overnight at 4°C with primary antibodies (see Table 2). After washing with PBST, blots were incubated with secondary antibodies for 1 hour at room temperature. The Odyssey imaging system (LICOR) was used for development, and analysis was performed using Image Studio Lite (LICOR). Beta-actin was used as a loading control for all samples.

### Open field

Locomotor activity was measured in a 45 × 45 × 80 cm open field (plywood, painted white) in a brightly lit room (70 lux). A ceiling-mounted camera recorded movement in the central and peripheral areas. After 30 minutes of habituation, each animal was placed in the center of the arena and recorded for 1 hour. Movement distance was analyzed using the EthoVision video-tracking software (version 16, Noldus, Netherlands) (Bonito-Oliva et al., 2016).

### Rotarod

The rotarod test (Ugo Basile, Gemonio, Italy, model 47600) evaluates motor coordination and learning. Mice were placed on a rotating rod above the instrument floor, adjusted to prevent injury but create a fear of falling. Time spent on the rod indicates motor skills, coordination, and balance. Using the Accelerarod protocol (4–40 rpm over 5 min), each trial ended when the mouse fell, completed a backward revolution, or reached 300 s. After 30 min habituation, mice underwent a 60 sec training trial at 4 rpm, followed by five test trials on day 1. On day 2, no habituation was given, and only one trial was performed. Each trial had a 5 min recovery period. Control animals generally show better performance between the first trial of day 1 and day 2 (Costa et al., 2004).

### Novel Home Cage

Each animal was singly housed in a new, unenriched home cage and allowed to habituate for 1 hour under bright light while being videorecorded. After habituation, the animals received an amphetamine injection (0.5–10 mg/kg, i.p., in saline) (Borgkvist et al., 2006) and were recorded for another hour. Locomotion was analyzed using EthoVision (version 16, Noldus, Wageningen, Netherlands). To confirm that the lack of locomotion at the highest dose (10 mg/kg) was due to severe stereotypies (Kelley, 2001; McNamara et al., 2006), we scored stereotypical movements according to a validated scale (Kelley, 2001).

### Patch-Clamp Electrophysiology

#### Brain slices preparation

Mice were euthanized with an overdose of pentobarbital, and 250μm coronal slices were obtained using a vibrating microtome (Leica VT1000S). Slices were prepared using a two-step NMDG-based protective recovery method (Lieberman et al., 2018; Ting et al., 2018). Initially, slices were made in an NMDG solution (in mM: 92 NMDG, 30 NaHCO3, 2.5 KCl, 20 HEPES, 2 thiourea, 1.25 NaH2PO4, 3 Na-pyruvate, 5 ascorbic acid, 25 glucose, 0.5 CaCl2, 10 MgCl2, pH 7.4). They were then stored in a HEPES-based holding solution (in mM: 92 NaCl, 30 NaHCO3, 2.5 KCl, 20 HEPES, 2 thiourea, 1.25 NaH2PO4, 3 Na-pyruvate, 5 ascorbic acid, 25 glucose, 2.5 CaCl2, 2 MgCl2, pH 7.4). All solutions were saturated with carbogen gas (95% O2, 5% CO2) throughout. After slicing, the tissue was stored in NMDG solution for 10 minutes at 32°C before being transferred to the HEPES solution.

#### Recording protocols

Patch-clamp recordings were conducted as previously described (Lieberman et al., 2018). Recordings were obtained using a patch clamp electrophysiology rig fitted with SciCam pro camera (Scientifica) and CoolLED pE-300ultra fluorescence illumination system, Multiclamp 700B amplifier (Molecular Devices) and Axon Digidata 1550B digitizer (Molecular Devices), using pCLAMP 11 software (Molecular Devices). Recording pipettes were fabricated from borosilicate glass microcapillaries (1.5mm diameter) and had resistances of 3–5 MΩ when filled with internal solution.

LTP was recorded in current-clamp configuration. Electrical stimulation was provided to the area slightly lateral and superficial to the recording area in layer 5/6. Subthreshold stimulation was given at a rate of 0.05Hz to establish a baseline response over 10 minutes. To potentiate the EPSP response, a theta-burst stimulation was given with 5 pulses and 2.5x current stimulation intensity, paired with postsynaptic depolarization to 0mV. The post-stimulation EPSP was recorded for 30 minutes after the initial stimulation at baseline stimulation intensity.

#### Analysis

All analysis of electrophysiological data was performed using the open-access Python-based software Stimfit 0.13.9 (Guzman, Schlögl, & Schmidt-Hieber, 2014). For LTP recordings, the average EPSP amplitude was calculated before and after stimulation, and the post-stimulation response was normalized to a baseline of 0%. A minimum threshold of 10% increase after 30 minutes was used to identify LTP.

#### Reagents

CTB488 (Cholera Toxin Subunit B, Alexa Fluor™ 488 Conjugate, #C22841) and Lucifer Yellow (#L453) were purchased from Invitrogen. SKF-38393 (1μM, #0922) and NKH-477 (10μM, #1603) were purchased from Tocris and dissolved in water. D-Amphetamine sulfate was purchased from Tocris and freshly prepared by dissolving in NaCl 0.9% solution (0.5-10 mg/Kg i.p., #2813, Tocris).

